# Effect and mechanism of black soybean peptides alleviating oxidative damage in the celiac disease cell model

**DOI:** 10.1101/2022.09.19.508472

**Authors:** Chenxu Cui, Na Wang, Enguang Gao, Xuefeng Sun, Qiuying Yu, Man Hu, Qian Xu, Ningnig Cui, Yuru Zheng, Chunfeng Wang, Fangyu Wang

## Abstract

Alpha gliadin peptide induces damage and apoptosis of intestinal cells and aggravates pathology of celiac disease (CD) by inducing oxidative stress. Therefore, inhibition or alleviation of oxidative stress in CD may be an effective approach to the adjunctive treatment of CD. Black soybean peptides (BSPs) have been shown to inhibit oxidative stress and inflammation. The effect of BSPs on CD remains unknown. In this paper, the effect and mechanism of BSPs on the α-gliadin peptide (p31-43)-induced Caco-2 cytotoxicity were studied. We identified BSPs that alleviated the cytotoxicity of p31-43 in the CD cell model: Caco-2 cells were pre-treated with bioactive peptides for 3 hours before the addition of p31-43 for treatment for 24 hours, and then cells were collected for subsequent experiments. Our results show that p31-43 can significantly increase the ROS and MDA levels of Caco-2 cells, disrupt the glutathione redox cycle, reduce the activity of the antioxidant enzyme, and inhibit the activation of antioxidant signaling pathways. BSPs pretreatment can inhibit the increase of Keap1 protein induced by p31-43, activate antioxidant genes through Nrf2 protein, improve the activity of the antioxidant enzyme, alleviates glutathione redox cycle imbalance, promote the expression of GCLC or GCLM, and reduce oxidative damage.

**Graphical Abstract:** Pattern of BSPs against oxidative damage in CD cell mode.

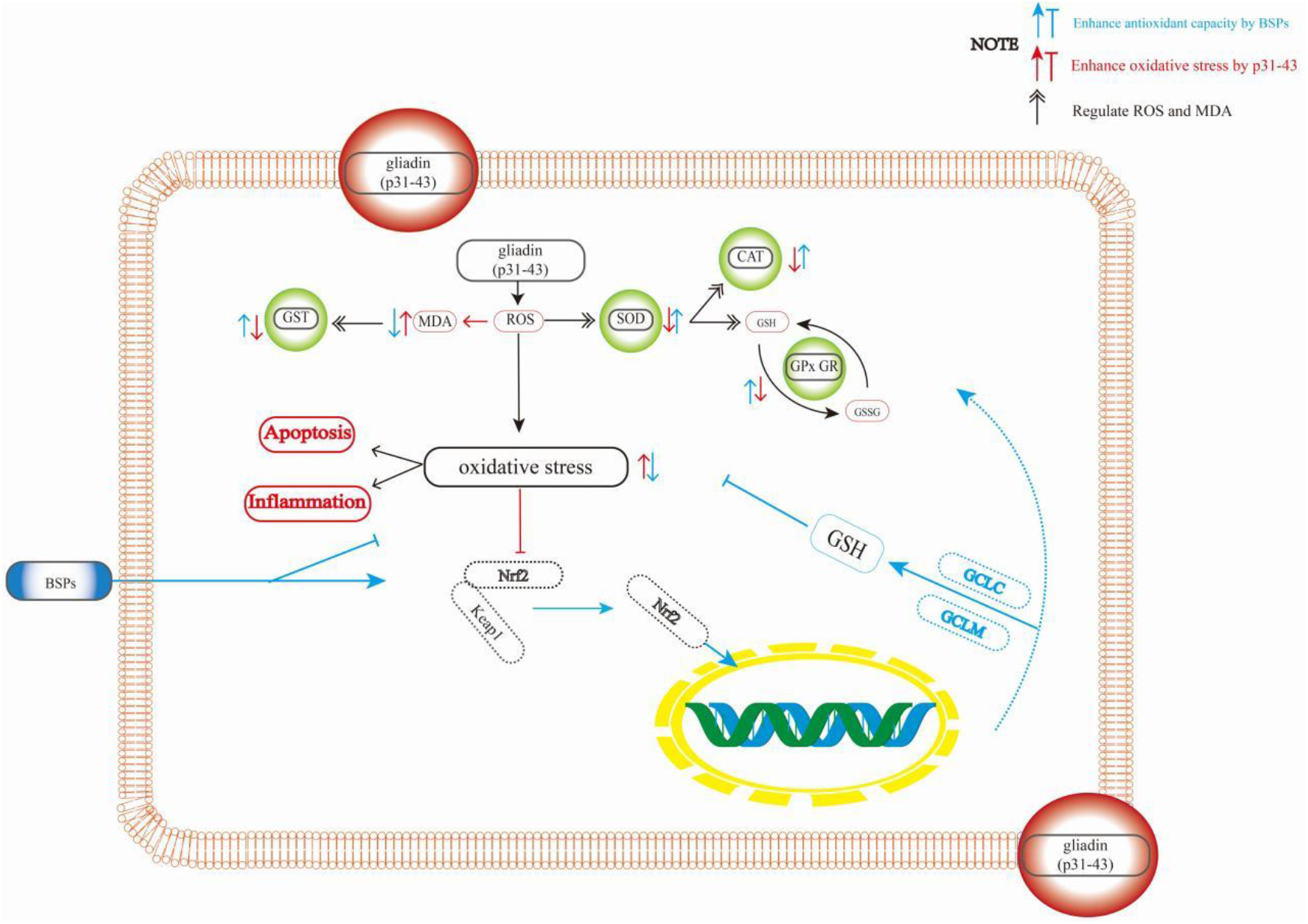

## 1. Introduction

CD is an allergic intestinal disease caused by genetically susceptible people (Rostami-Nejad et al., 2014) (carrying HLA-DQ2 or HLA-DQ8 genes) ingesting gluten-containing wheat and other wheat-cereal crops. The current worldwide incidence of CD is 1.4% (A et al., 2018), and the prevalence of CD has increased dramatically and has become a serious chronic gastrointestinal disorder in Europe. Gluten proteins are mainly composed of gliadin and glutenin, which can’t be completely hydrolyzed in the intestinal lumen to produce different peptides. They trigger the adaptive immune response mediated by CD4^+^Th1 cells (Bayardo et al., 2012) and the innate immune response mediated by intraepithelial lymphocytes (Barone et al., 2007; Luigi et al., 2003), respectively, and jointly cause intestinal inflammation, resulting in lamina propria and epithelial inflammatory cells infiltrating villus atrophy and crypt hyperplasia.

Oxidative stress, inflammatory response, and intestinal lesions played important roles in CD (Dheen et al., 2007), Gluten enhanced the t-cell inflammatory response through oxidative stress, are responsible for cell death via oxidation of DNA, proteins, lipids, and almost any other cellular constituent, and induced an immune response to TGM2. Gluten (Tarique et al., 2016) also promoted the expression of COX-2, iNOS, and inflammatory mediators such as NO, PGE2, IFN-γ, TNF-α, IL-1β, IL-6, and IL-15 by increasing the level of ROS (Maiuri et al., 2005), induced inhibition of PPAR-γ and activating MAPK, NF-κB signaling pathways(Caputo et al., 2017; Maiuri et al., 2005; Patlevič et al., 2016; Saskia et al., 2016), leading to the enhanced inflammatory response in the body. Oxidative stress and inflammation disrupt cell membrane integrity, disrupt intestinal permeability, and enhance environmentally induced toxicity, resulting in intestinal cell injury and apoptosis, cell proliferation, and intestinal epithelial barrier and function (Qin and Hou, 2016).

Therefore, ROS has been considered one of the major mechanisms involved in CD pathogenesis and progression, which affects cell morphology, proliferation, and viability (R et al., 1999; Sabatino et al., 2001). Oxidant stress is caused by an imbalance between reactions and responses to ROS and antioxidant defense systems. ROS include superoxide radicals (O_2_^•-^), hydroxyl radicals (•OH) and hydrogen peroxide (H_2_O_2_), and singlet oxygen (SATO et al., 2013). There are several antioxidant defense mechanisms in biological systems that regulate ROS production, but studies have shown that several antioxidant defense mechanisms get damaged in people with CD.

Biological systems play an important role through enzymatic and non-enzymatic cellular antioxidant systems to remove excess accumulated ROS. SOD-CAT reduces oxidative stress by decomposing ROS into H_2_O_2_ and O_2_ (Aksoy et al., 2004; Chakravarti et al., 2015; Goyal et al., 2012; Patlevič et al., 2016; SATO et al., 2013). But Zhang (Vesnać et al., 2012) found that in the CD cell model, gliadin inhibited the activity of Caco-2 cells, significantly increased the levels of ROS and MDA, and decreased the level of SOD activity. Excessive ROS can also be removed by the GSH pathway under the catalysis of the GPx enzyme (Arthur, 2000). GSH is a powerful antioxidant, antitoxin, and enzyme-cofactor (Berkholz et al., 2008; Outten and Culotta, 2004), GPX catalyzes the reaction of GSH with ROS to generate H2O2 and oxidized glutathione (GSSG). GR is an equally important antioxidant enzyme (Grès et al., 1998; Ribeiro et al., 2012), which plays a critical role in GSH regeneration: GR catalyzes the generation of GSH from GSSG through an NADPH-dependent mechanism. The interconversion of GSH and GSSG is called the glutathione redox cycle, it is an important cycle of excessive ROS removal. But studies (Vesnać et al., 2012) have shown that in patients with CD, a significant malfunction of the glutathione redox cycle with a concomitant decrease in the capacity to regenerate GSH. In another in vitro study, Elli (L et al., 2003) found that gliadin caused a decrease in glutathione content and decreased GR, GPx, and GST activities. GST (Shi et al., 2014) catalyzes GSH to detoxify peroxidation lipid by-products. In addition, the primary treatment for CD is lifelong adherence to a gluten-free diet (GFD), but the glutathione redox cycle was found to be impaired in patients who adhered to GFD. Because of key mechanisms of oxidative stress response in CD, it is worth further exploring whether antioxidant substances have the potential to regulate or alleviate CD.

Current studies suggest that Nrf2 is an important signaling pathway that activates the cellular defense system against oxidative damage and inflammation. Nrf2 generally binds to Keap1 in the cytoplasm of normal cells (Itoh et al., 1999), and Keap1 degrades Nrf2 to balance Nrf2 protein expression. However, under the stimulation of other signals such as oxidative stress, Nrf2 is activated after separation from Keap1 to activate Nrf2-targeted antioxidant genes, it includes heme oxygenase 1 (HO-1), NAD(P)H, GCL, SOD, CAT, GR, GPx, GST, and maintains the oxidative/antioxidant balance in cells by up-regulating their expression (Venugopal and Jaiswal, 1998). GCL is a heterodimeric protein composed of a catalytic subunit (GCLC) and a regulatory subunit (GCLM), which catalyzes the synthesis of GSH and plays a key role in antioxidant defense (Jiang et al.).

The protein content of black soybean can reach more than 40%–50% and the type and content of amino acids are in line with the proportion of nutrition required by the human body. And studies have found that antioxidant active functional peptides from black bean protein hydrolysates, can inhibit the generation and accumulation of free radicals in the body and reduce the level of ROS. As a plant-derived active peptide, great progress has been made in the study of its anti-inflammatory and antioxidant activities in vitro. For example, black bean peptide can reduce the content of MDA in serum and liver, and significantly increase the activity of SOD in liver and GPx in serum and liver (Singh et al., 2014).

Therefore, this study will establish a cell model of CD based on the p31-43 induced Caco-2 cells to explore: (1) Mechanism of oxidative damage in CD cell model. (2) Mechanism of black bean peptide alleviating oxidative damage in CD cell model.

## 2. Materials and methods

### 2.1. Reagents and antibodies

High-glucose Dulbecco’s modified Eagle’s medium (DMEM), fetal bovine serum (FBS), ROS assay kit, MDA assay kit, SOD assay kit, GSH/GSSG assay kit, CAT assay kit, GST assay kit, GR assay kit, GPx assay kit were purchased from Beyotime (Shanghai, China) and Solarbio (Beijing, China). The Cell Counting Kit-8 (CCK-8) was purchased from Meilunbio (Dalian, China). Primary antibodies against, β-actin, and Nrf2 were purchased from Cell Signaling Teart (Shanghai, China). Conjugated secondary antibodies (anti-rabbit and anti-mouse) were purchased from Proteintech (Wuhan, China). All other chemicals were of analytical grade and purchased from Solarbio (Beijing, China) and Beyotime (Shanghai, China) unless otherwise stated.

### 2.2. Cell culture and treatment

Caco-2 cell line derived from human Colorectal Adenocarcinoma was used as a model of human intestinal epithelial cells and has been widely used in vitro studies of coeliac disease injury. Caco-2 cells (ATCC, Manassas, VA) were cultured in DMEM supplemented with 10% (v/v) FBS, 1% (v/v) penicillin and 1% (v/v) streptomycin (Gibco) at *37°C* in a humidified 5% CO_2_ atmosphere. Subculture cells when they are about 80% confluent, cells were subcultured every 3 days at a subcultivation ratio of 1:3 by trypsinization.

Treatment of Caco-2 cells according to experimental requirements. p31-43 and bioactive peptide were dissolved in DMEM and diluted to 100 μM, and Vc was dissolved in DMEM and diluted to 250 μM. Normal control: Caco-2 cells were cultured in a cell culture plate without treatment. The positive control group: Caco-2 cells were cultured in a cell culture plate for 24h and treated with 250μM Vc for 3h. The bioactive peptide group: Caco-2 cells were cultured in a cell culture plate for 24h and treated with 100μM bioactive peptide for 3h.

The CD model group: Caco-2 cells were cultured in a cell culture plate for 24h, then treated with PBS for 3h before treatment with p31-43 for 24 h. The BSPs pretreatment CD model group: Caco-2 cells were cultured in a cell culture plate for 24h and treated with 100μM BSP for 3h before treatment with p31-43 for 24 h. The positive control group: Caco-2 cells were cultured in a cell culture plate for 24h and treated with 250 Vc for 3h before treatment with p31-43 for 24 h.

### 2.3. Caco-2 Cell Viability Assay

According to different experimental requirements, after the cells are treated, cells were cultured in a complete culture medium containing 10% (*v*/*v*) CCK-8 solution for 2 h and the absorption was recorded at 450nm by using a microplate reader (POLARstar Omega, BMG LABTECH) according to the manufacturer’s protocol.

### 2.4. Flow cytometry detection

According to the manufacturer’s protocol, collected cells after different treatments, centrifuged for 5 min (1000 r/min), and discarded supernatant, the sample solution was incubated with annexin V-FITC for 20 min at room temperature in the dark, then added the binding buffer to sample tube. The samples were analyzed using a CytoFLEX flow cytometer (Beckman Coulter, Brea, CA, USA).

### 2.5. Determination of ROS levels

ROS levels of the normal cell group, the CD cell model group, the bioactive peptide group, and the BSPs pretreatment CD model group were determined respectively by using a microplate system according to the manufacturer’s protocols of ROS assay Kit.

Caco-2 cells were cultured in 96-well plates (1×10^4^ cells per well) according to the experimental requirements, after culture, the cells were carefully rinsed with PBS once, then add 100μL DCFH-DA probe (10μM) and incubated for 1 h. Then, cells were carefully washed 3 times by an incomplete medium to remove the DCFH-DA probe. Last, the fluorescence intensity was measured at the excitation wavelength of 488 nm and emission wavelength of 525 nm.

### 2.6. Determination of MDA levels

MDA levels in different groups were detected. Caco-2 cells were treated in 6-well plates (2×10^5^ cells per well) according to the experimental requirements. Caco-2 cells were washed with PBS, then cells were collected and lysed by Cell lysis buffer containing protease and phosphatase inhibitor cocktail on ice. The cell Lysis buffer was centrifuged (12000 r/min) for 15 min. Subsequently, the values were determined with a microplate reader and the MDA content was calculated.

### 2.7. Measurement of GSH and GSSG levels

Detected GSH and GSSG levels in different groups. After the Caco-2 cells were cultured in 12-well plates (2×10^5^ cells per well) according to the experimental requirements, washed cells with ice-cold PBS, then the cells were collected by centrifugation and the supernatant was discarded, finally, the GSH and GSSG contents of cells were detected and calculated according to the detection kit.

### 2.8. Measurement of antioxidant enzymes

Caco-2 cells were cultured in 12-well plates (2×10^5^ cells per well) according to the experimental requirements. After culture, the cells were treated according to the detection protocol of the detection kit, and the antioxidant enzyme activities of the cells were measured and calculated.

### 2.9. Cellular Western blot analysis

Caco-2 cells were treated according to experimental requirements and then Caco-2 cells were washed with PBS and lysed in RIPA Lysis buffer containing protease and phosphatase inhibitor cocktail on ice for 10 min. Subsequently, the cell lysis buffer was centrifuged under 12000 rpm for 15 min (4°C), last, the supernatant was added protein loading buffer to denature by heating at 100°C for 10 min.

Protein samples were transferred to polyvinylidene fluoride (PVDF) membrane by 12.5% sodium dodecyl sulfate-polyacrylamide gel electrophoresis (SDS-PAGE). The membrane was blocked with 5% bovine serum albumin (BSA) in TBST or 5% skim milk in PBST for 2 h at room temperature and then incubated overnight at 4°C with the indicated primary antibodies. Subsequently, the membrane was washed six times for six min using TBST / PBST and incubated for 1h with the secondary antibody. Finally, the washes were repeated. Protein signaling was shown using an enhanced chemiluminescence kit and the ChemiDoc XRS imaging system.

### 2.10. Statistical analysis

All experiments were repeated at least three times. SPSS software was used to analyze data, Image J software was used to analyze the Western blot data. Measurements were expressed as the mean ± SD. Differences among multiple groups were compared using one-way ANOVA. Differences between pairs of groups were assessed using Tukey’s test for multiple comparisons. The differences were considered significant at the *P* < 0.05

## 3 Results and discussion

### 3.1. Cell viability

Cell viability and apoptosis are often used as an indicator of cytotoxicity.

Detected the activity of Caco-2 cells to determine whether the bioactive peptides (100μM) had cytotoxicity. CCK-8 results (Figure 1) show that 100μM of bioactive peptides did not have adverse effects on Caco-2 cell viability (cells viability: ≥ 95%). It is proved that bioactive peptides can be used in the following experiments.

**Figure 1.**
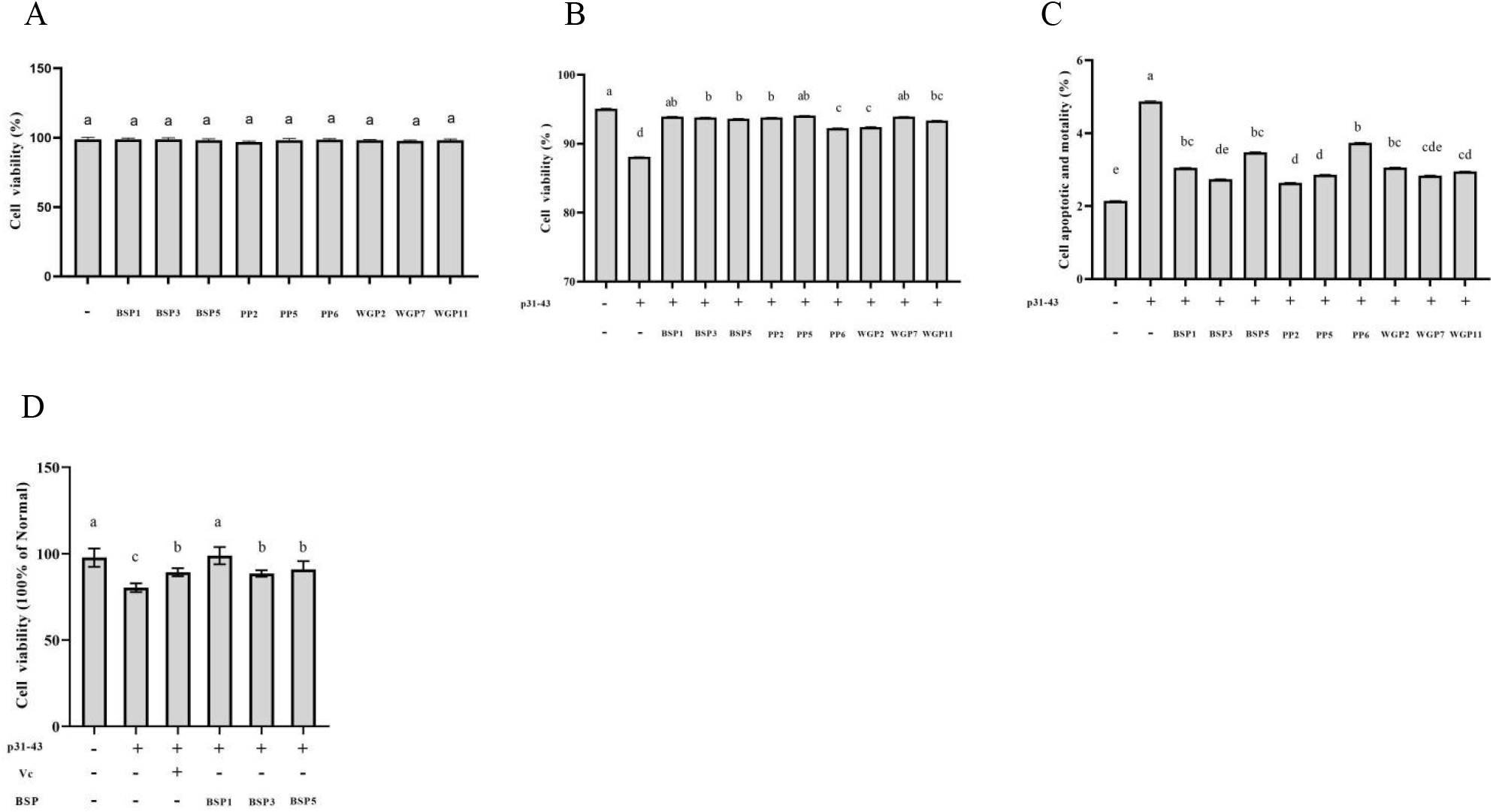
Effects of bioactive peptide in Caco-2 cells viability. (A) Toxic effect of the bioactive peptide on Caco-2 cells was measured using the CCK-8 assay. (B) Effects of the bioactive peptide on the viability of P31-43-induced Caco-2 cells obtained by flow cytometry. (C) Effects of the bioactive peptide on the Cell apoptotic and mortality of P31-43-induced Caco-2 cells obtained by flow cytometry. (D) The toxic effect of BSPs in the CD cell model was measured using the CCK-8 assay. Different letters represent significant differences. Statistical significance and highly significant were set at *P* < 0.05 and *P* < 0.001.

Flow cytometry results (Figure 1,2) showed that Caco-2 cells exposed to p31-43 significantly decreased cell viability and increased apoptosis and mortality, (*P* <0.001), suggesting that Caco-2 cells were injured by p31-43. Compared with the CD model group, the cell viability and apoptosis were found to be improved significantly (*P* < 0.05) in the bioactive peptides pretreatment CD model group, this indicates that various bioactive peptides had protective effects. In the present study, BSPs were selected as the research object for the follow-up study. In order to confirm whether BSPs could alleviate the decrease of cell activity in the CD cell model by CCK-8 assay kit. CCK-8 results (Figure 1) show that p31-43 induced a decrease in cell viability in the CD cells model, and pretreatment of BSPs could improve the activity of the CD cells model.

**Figure 2.**
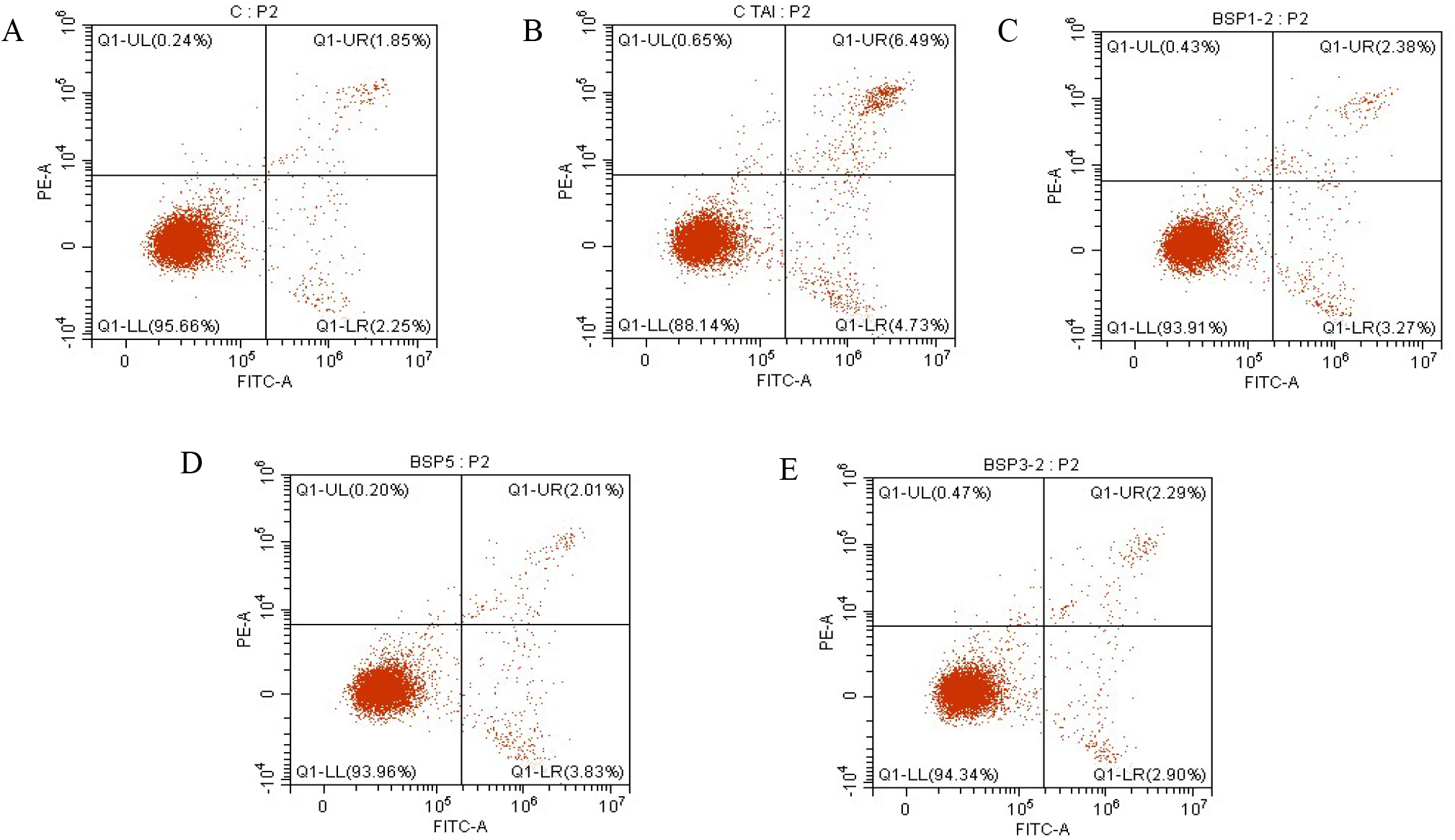
Flow cytometry results. (A) normal cells. (B) P31-43-induced Caco-2 cells. (C)-(E) BSP3 pretreated CD cell model.

These results demonstrate that p31-43 can induce cell damage, and bioactive peptides can indeed alleviate p31-43-induced cytotoxicity. These results are in agreement with Kramer’s view (Kramer et al., 2019). The p31-43 and p31-49 of gliadin were found to enter cells by endocytosis, also increase levels of ROS, cause oxidative stress (Kramer et al., 2019; Zimmer et al., 2010), and decrease cell viability. Excessive ROS not only destroys mitochondrial integrity, increases the expression of apoptotic proteins, and leads to cell apoptosis, but also destroys nucleic acid and protein structures and cell membrane structures to lead to cell death (Angélica et al., 2019; Cremonini et al., 2017; Iglesias et al., 2020; Maren et al., 2008; Shaojun et al., 2020). Therefore, we infer that p31-43 induced oxidative stress in the CD model. Experimental results show that BSPs can alleviate cell damage in the CD model, which may be related to its antioxidant capacity because some scholars have found that antioxidants can inhibit the inflammatory response of the Caco-2 cell model induced by gliadin and reduce the cell membrane damage in CD cell model. It also regulates the intestinal barrier by inhibiting oxidative stress and inflammation-related signaling pathways (Zhiyan et al., 2011). Therefore, we tested our hypothesis by measuring oxidative stress indicators.

### 3.2. Preventive effects of BSPs on ROS and MDA generation

ROS and MDA are great measures of oxidative stress. MDA is one of the complex compounds produced by the oxidation of some fatty acids when cells are subjected to oxidative stress. It was earlier shown that excessive ROS could damage cell membrane integrity by oxidizing and degrading unsaturated fatty acids (lipid peroxidation) in cell membranes. lipid peroxidation was involved in the cleavage of polyunsaturated fatty acids at its double bonds, resulting in cellular membrane damage and an increase in MDA production. Previous studies have shown that gliadin can significantly increase ROS levels (Van Buiten et al., 2018), which is also confirmed by our experimental results, p31-43 extremely significantly (*P* <0.01) increased the level of ROS and MDA in cells (Figure 3). It was proved that exposure of Caco-2 cells to p31-43 induced peroxidation stress.

**Figure 3.**
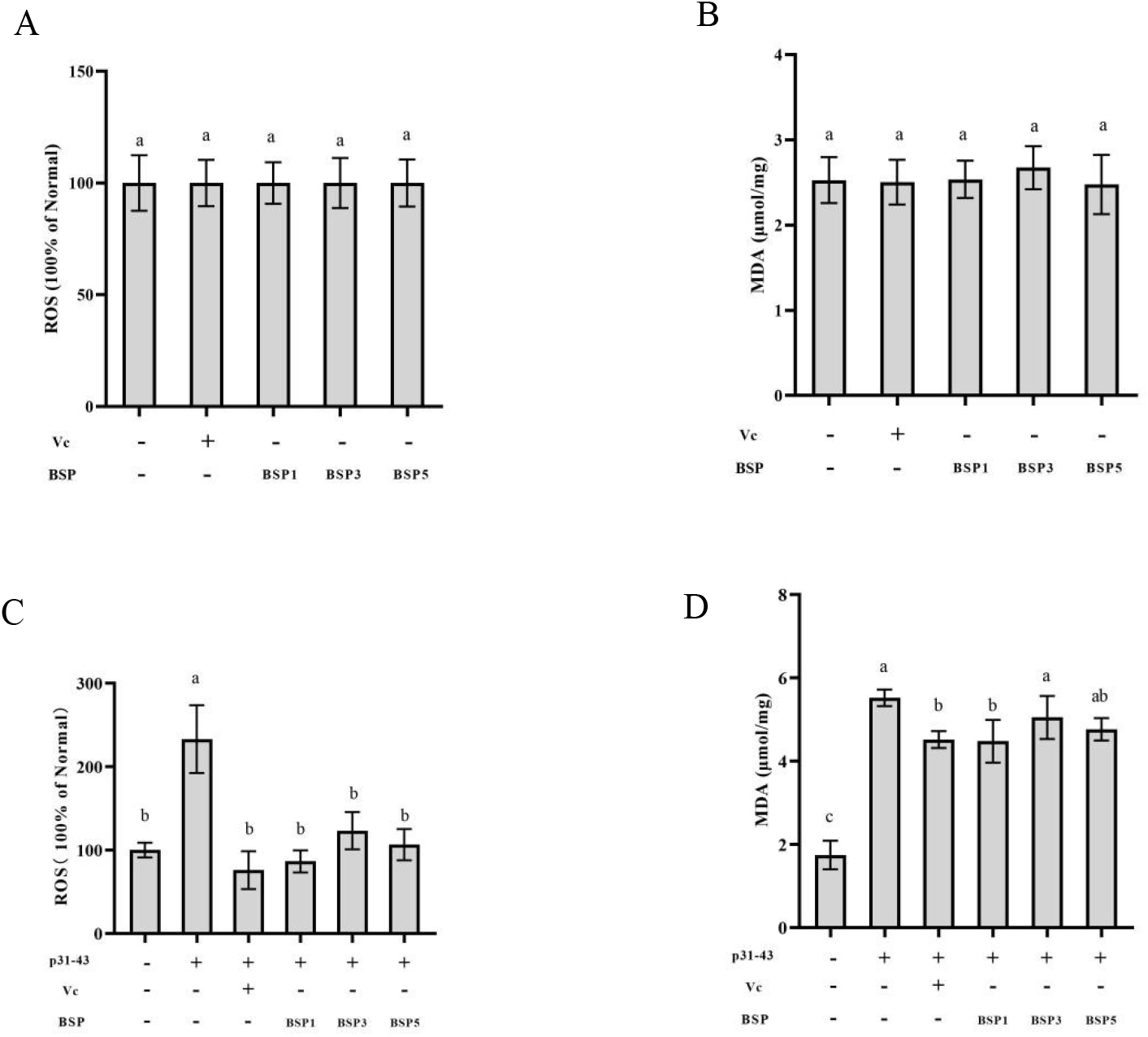
Effect of BSPs on oxidative stress. (A) Effect of BSPs on ROS in normal cells. (B) Effect of BSPs on MDA in normal cells. (C) Effect of BSPs on ROS in CD cell model. (D) Effect of BSPs on MDA in CD cell model. Different letters represent significant differences. Statistical significance and highly significant were set at P < 0.05 and P < 0.001.

Treatment of BSPs had no effect on levels of ROS and MDA in normal cells but compared with the CD group, BSPs pretreatment reduced significantly the levels of ROS, and BSP1 could also significantly reduce the level of MDA. The results showed that BSP pretreatment may inhibit oxidative stress or improve the antioxidant capacity of cells in the CD model. Different BSPs exhibited the ability to reduce ROS on HepG2 cells, THP-1 macrophages, and Caco-2 (Lule et al., 2015; Wenyi). The antioxidant capacity of different types of BSBs on different cells was not identical, which may be related to their intrinsic mechanism of action, these may be the reasons why pretreatment of BSP3 and BSP5 didn’t reduce MDA content in the CD model. Studies found that peptides can inhibit lipid oxidation through multiple pathways including inactivation of reactive oxygen species, scavenging free radicals, enhancing or protecting antioxidant enzyme activity, and so on (Singh et al., 2014). Therefore, this study explored the mechanism of black soybean peptide against oxidative damage in the CD cell model from the changes in glutathione redox cycle, antioxidant enzyme activity, and antioxidant signaling pathway.

### 3.3. Preventive effects of BSPs on glutathione redox cycle in CD cells model

Guo (Yi et al., 2020) proved that some soybean peptides can regulate oxidative stress of cells by measuring the content changes of GSH and GSSG. In order to further study the principle of BSPs against oxidative stress in the CD cells model, we studied the effect of BSPs on GSH and GSSG. GSH plays an important role in the detoxification of oxidative stress, and the optimal balance of the GSH to GSSG ratio is essential for cell survival. GSH to GSSG ratio has been considered an indicator of oxidative stress (Dheen et al., 2007).

Treatment of BSPs did not affect the glutathione redox cycle in normal cells (Figure 4). In the CD cells model, p31-43 disrupted the glutathione redox cycle, which was characterized by downregulation of GSH and a significant increase (*P* < 0.05) in GSSG levels (Figures 4), and a significant decrease (*P* < 0.05) in the ratio of GSH to GSSG. As shown in the result, pretreatment of BSP1 significantly increased the level of GSH (*P* < 0.05), meanwhile, a decline in GSSG was observed in the pretreatment of the BSPs group (*P* < 0.05).

**Figure 4.**
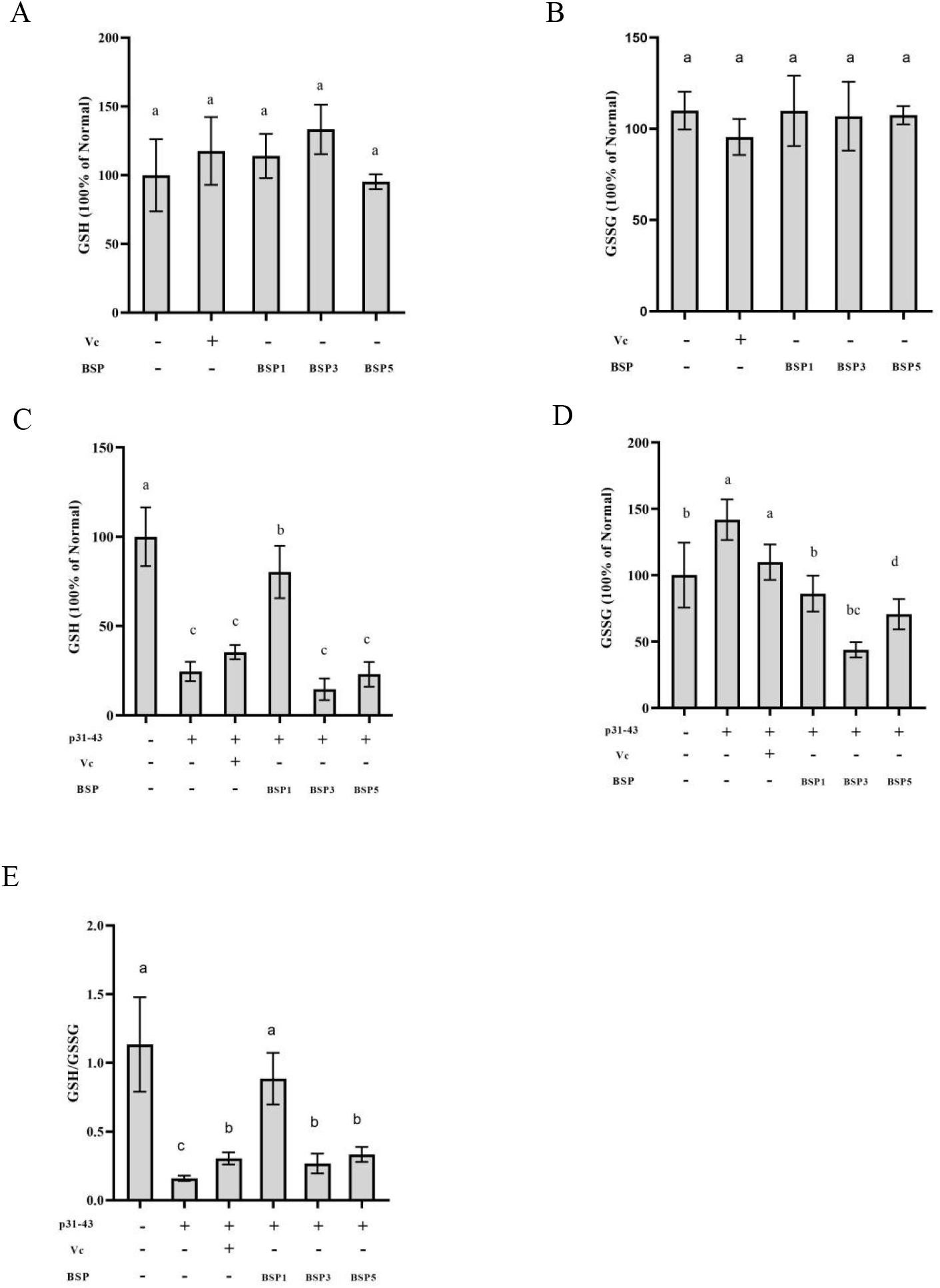
Effect of BSPs on GSH and GSSG levels in cells. (A) Effect of BSPs on GSH in normal cells. (B) Effect of BSPs on GSSG in normal cells. (C) Effect of BSPs on GSH in CD cell model. (D) Effect of BSPs on GSSG in CD cell model. (E) Effect of BSPs on GSH/GSSG in CD cell model. Different letters represent significant differences. Statistical significance and highly significant were set at P < 0.05 and P < 0.001.

According to studies (Chen et al., 2016; Eun?Young et al., 2018; Guo et al., 2018), we know that GSH is catalyzed by GSSG during oxidation, so GSH consumption and GSSG production reflect the degree of intracellular oxidation. Therefore, the results proved that p31-43 decreased the levels of GSH and increased the levels of GSSG, disrupting the balance of glutathione/glutathione ratio to induced oxidative in the CD cells model, this phenomenon also occurs in patients with CD (Vesnać et al., 2012) and other models of oxidative stress (Patlevič et al., 2016; Yi et al., 2020). Studies (Yi et al., 2017) showed that treatment of antioxidant peptides increased the content of GSH compared to the oxidative stress group. Consequently, we concluded that BSP1 protects against oxidative stress by increasing or protecting GSH content.

### 3.4. Preventive effects of BSPs on the activities of antioxidant enzymes in the CD model

Previous experiments have found that both CD cell models and peptide antioxidant activity are associated with the glutathione redox cycle which plays an important role in the cellular antioxidant defense system, and the related antioxidant enzymes are also important. To explain the role of the glutathione redox cycle in the CD model, we learn from researches (Goyal et al., 2012; Yi et al., 2020) show that the antioxidant enzymes SOD, CAT, GPx, GR, and GST were inhibited in Preventive effects. So, we studied the effect of BSPs on antioxidant enzymes.

SOD and CAT are the main antioxidant enzymes in the human body, and they play an important role in the defense mechanism against oxidative damage. The treatment of BSP5 (Figure 5) significantly increases the activity of SOD in normal cells, and treatment of BSPs could significantly increase (*P* < 0.05) the activity of CAT (Figure 5). However, pretreatment of BSPs didn’t protect the activity of SOD, and pretreatment of BSP1, BSP5, and Vc promoted (*P* < 0.05) activity of CAT in the CD cell model (Figure 5). The results showed that p31-43 inhibited (*P* < 0.05) the activity of antioxidant SOD, which may be due to the accumulation of ROS leading to the damage of intracellular biomolecules, thus changing the enzyme activity, and other models of oxidative stress showed a similar pattern (Zhu et al., 2013), this may also be the reason why BSP5 increased activity of SOD in Caco-2 but didn’t protect SOD activity in CD cells model. The decrease of activity of SOD also led to a decrease in the efficiency of eliminating ROS, forming a vicious circle. It is consistent with other conclusions about oxidative stress (Vivek and Saba, 2020; Zhu et al., 2013). According to the results, we concluded that one of the ways for BSP1 and BSP5 to inhibit the oxidative damage of p31-43 is to improve the antioxidant capacity of cells by enhancing and protecting CAT activity.

**Figure 5.**
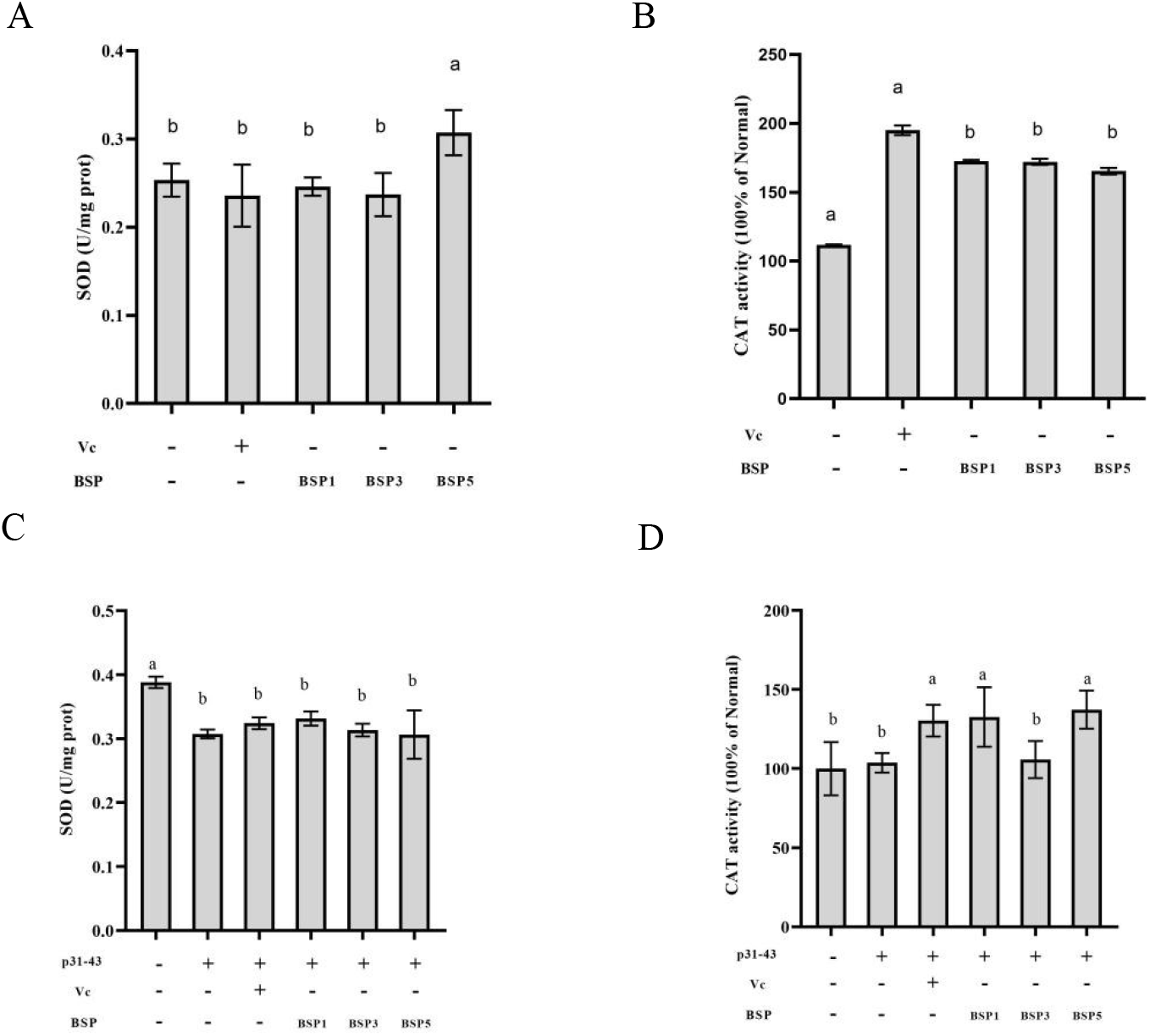
Effect of BSPs on the activity of antioxidant enzymes in cells. (A) Effect of SOD on GSH in normal cells. (B) Effect of CAT on GSSG in normal cells. (C) Effect of BSPs on SOD in CD cell model. (D) Effect of BSPs on CAT in CD cell model. Different letters represent significant differences. Statistical significance and highly significant were set at P < 0.05 and P < 0.001.

The glutathione redox cycle is an important cycle of excessive ROS removal, so the related antioxidant enzyme activity also plays an important role in the cellular antioxidant defense system, so the effect of BSPs on GR, GPx, and GST should be explored.

The contents of GR, GPx, GST, and other antioxidant enzymes in the cells of each experimental group were determined, and the results were shown in Figure 6. The treatment of BSP1 and BSP3 significantly enhanced (*P* < 0.05) the activity of GR in normal cells, but only pretreatment of BSP1 and Vc significantly increased (*P* < 0.05) the activity of GR in the CD cell model. For GPx, Vc and BSP5 significantly inhibited the activity of GPx in normal cells, resulting in their pretreatment could not alleviate the decline of GPx activity in the CD model, and even exacerbated the injury, but BSP1 could significantly improve the activity of GPx in the CD model (*P*<0.05). Compared with the normal group, BSP3 decreased the activity of GST in normal cells. However, compared with the CD cells model, only the pretreatment of BSP1 significantly protected the activity of GST.

**Figure 6.**
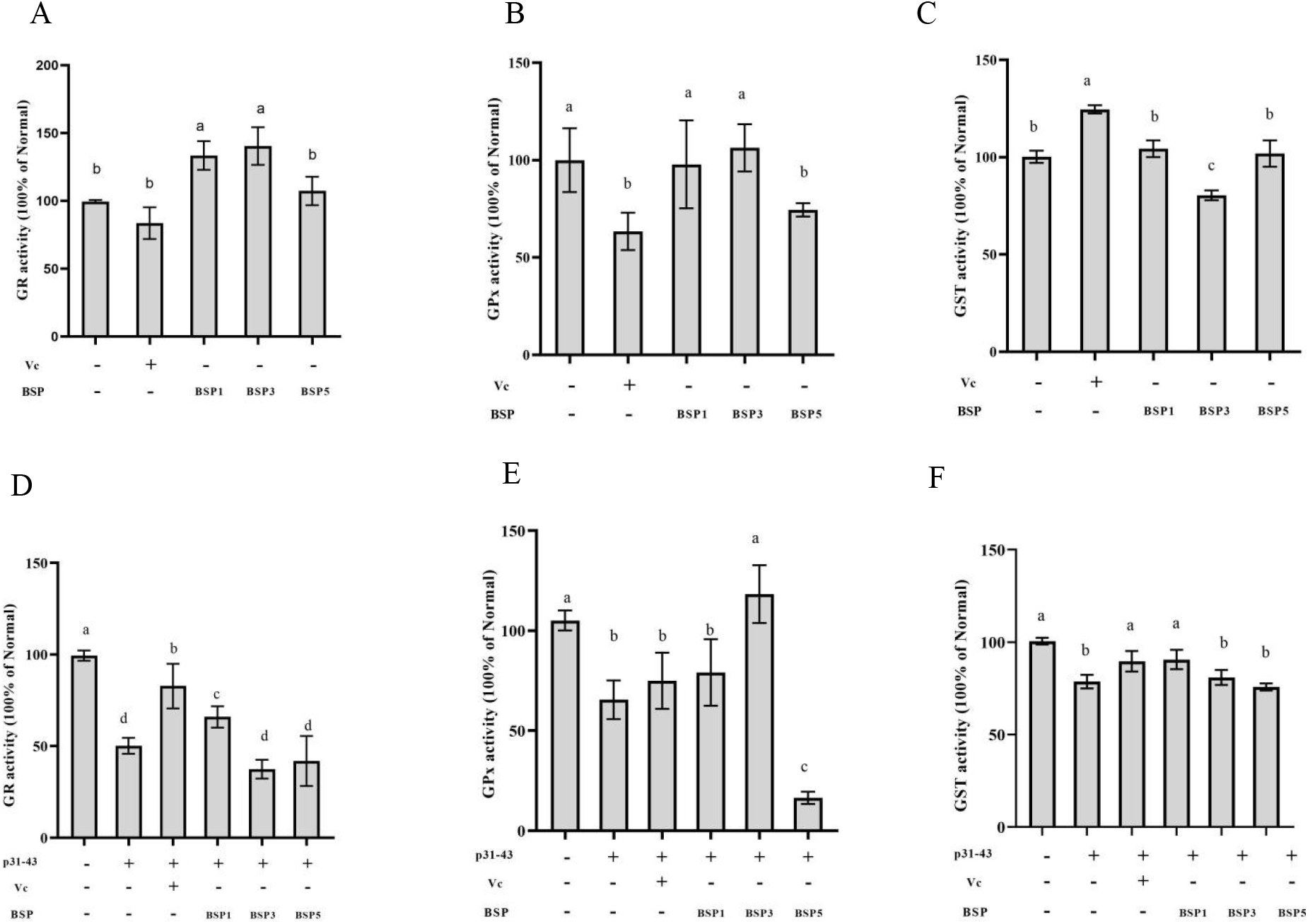
Effect of BSPs on the activity of antioxidant enzymes in cells. (A) Effect of GR on GSH in normal cells. (B) Effect of GPx on GSSG in normal cells. (C) Effect of GST on GSSG in normal cells. (D) Effect of GR on GSH in CD cells model. (E) Effect of GPx on GSSG in CD cells model. (F) Effect of GST on GSSG in CD cells model. Different letters represent significant differences. Statistical significance and highly significant were set at P < 0.05 and P < 0.001.

Based on the results, we infer that p31-43 might reduce the activities of GR, GPX, and GST enzymes in the CD model, resulting in the inhibition of GSH scavenging ROS and GST detoxification of lipid peroxides, leading to an increase in the degree of cellular oxidative stress.

The result has shown that BSPs had different effects on the activities of GR, GPx, and GST.

BSP1 can promote the activity of GR and protect GR from the inhibition of the p31-43 peptide, by this way, the GSSG produced by cells alleviating p31-43-induced ROS could rapidly become GSH, which is used again to resist oxidative stress; This also resulted that the content of GSH in BSP1 pretreatment group was significantly higher than that in CD model, and the content of GSSG was lower than that in the p31-43 group, which led to the recovery of GSH/GSSG ratio close to the normal cell level. Studies (Shi et al., 2014; Vivek and Saba, 2020; Zhu et al., 2013) have also found that antioxidants can enhance the ability of cells to resist oxidative stress by increasing the activity of GR enzymes. We also found that although BSP1 did not promote the expression of the GST enzyme, it alleviated the inhibitory effect of the p31-43 peptide on the GST enzyme, resulting in a significant decrease in MDA content in the CD model group pretreated with BSP1.

BSP3 can not only promote the activity of GR in normal cells but also protect the activity of GPx in the CD model. As a result, GPx can catalyze the reaction of GSH with excessive ROS to produce GSSG, while GR catalyzes the regeneration of GSH, but with the stimulation of p31-43, the activity of GR was also inhibited, which leads to the decrease of GSH regeneration ability, and ultimately leads to the decrease of the content of GSH and GSSG. However, BSP3 inhibited the activity of GR in normal cells, resulting in no significant difference in MDA level between the BSP3 pretreated group and the CD model group.

BSP5 pretreatment did not promote and protect GR, GPx, and GST in the CD model, indicating that BSP5 pretreatment alleviated oxidative stress mainly through SOD and CAT pathways, which also explained the reason why MDA level was not alleviated in the BSP5-pretreated group.

In this study, we found that some BSPs could increase the activity of antioxidant enzymes in normal cells, which were still inhibited after p31-43 stimulation, this may be related to the inhibition of antioxidant defense molecules such as Nrf2. Therefore, we will study the mechanism of BSPs alleviating oxidative damage in the CD model at the molecular level.

### 3.5. Preventive effects of BSPs on Keap1/Nrf2 sign pathway

Studies have shown that Nrf2 is involved in ROS regulation in vivo, and the antioxidant function of most antioxidants is usually related to the activation of the Nrf2 signaling pathway (Vivek and Saba, 2020).

In normal cells, BSPs could promote the expression of Nrf2 protein but did not affect the expression of other proteins (Figure 7).

**Figure 7.**
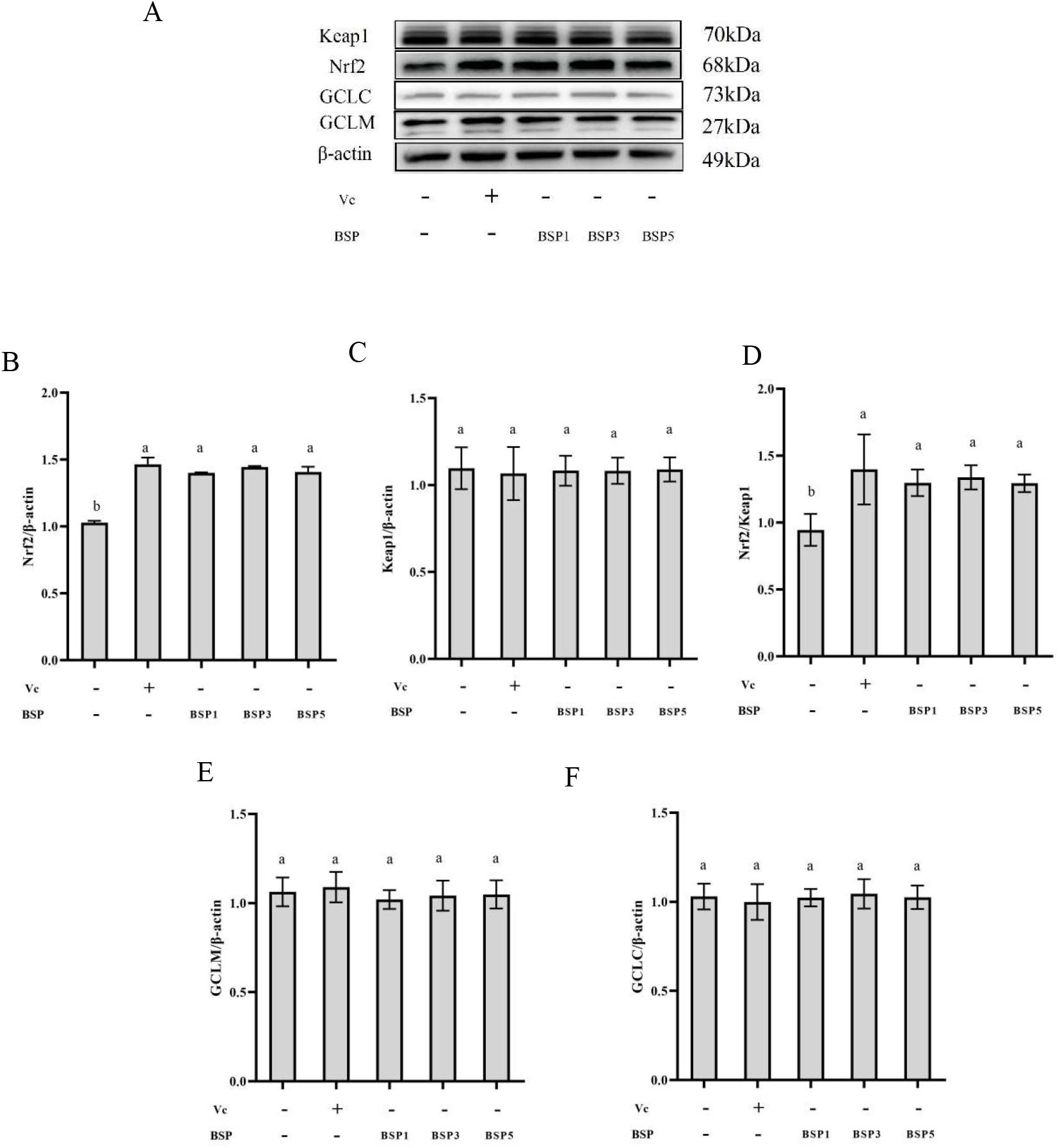
Effects of BSPs on the expression of antioxidation signaling pathway-related proteins in normal cell. (A) Western blot analysis of Keap1, Nrf2, GCLC, GCLM, and β-actin in a normal cell. (B) Quantitative analysis of Keap1 levels. (C) Quantitative analysis of Nrf2 levels. (D) Quantitative analysis of GCLC levels. (E) Quantitative analysis of GCLM levels. (F) Quantitative analysis of β-actin levels. Different letters represent significant differences. Statistical significance and highly significant were set at *P* < 0.05 and *P* < 0.001.

Compared with the normal group, p31-43 affected (*P* < 0.05) Nrf2/Keap1 and GCLC signaling pathways (Figure 8). The expression of Keap1 protein and Nrf2 protein (Figure 8) in the CD model were significantly increased (*P* < 0.05), but the value of Nrf2/Keap1 was significantly decreased (*P* < 0.05), indicating that the Nrf2 antioxidant pathway in the CD model group was inhibited. The expression of GCLC in the CD model was significantly lower than that in the normal group (Figure 8), but there was no significant difference in GCLM protein expression, indicating that p31-43 could inhibit the expression of GCLC protein and had no significant effect on GCLM.

**Figure 8.**
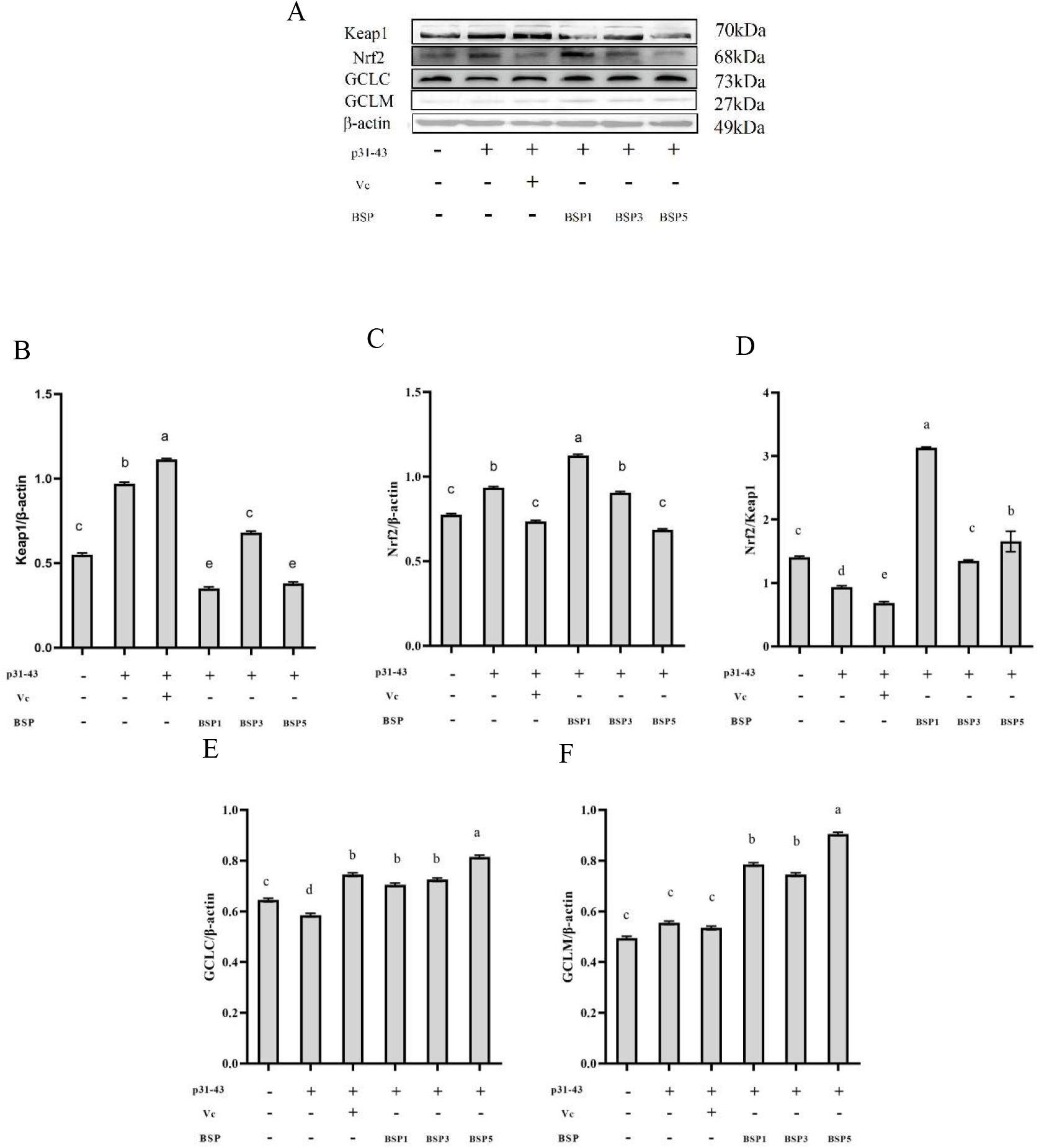
Effects of BSPs on the expression of antioxidation signaling pathway-related proteins in CD cell model. (A) Western blot analysis of Keapl, Nrf2, GCLC, GCLM, and β-actin in CD cell model. (B) Quantitative analysis of Keapl levels. (C) Quantitative analysis of Nrf2 levels. (D) Quantitative analysis of GCLC levels. (E) Quantitative analysis of GCLM levels. (F) Quantitative analysis of β-actin levels. Different letters represent significant differences. Statistical significance and highly significant were set at P < 0.05 and P < 0.001.

The results showed that the expression of Nrf2 and Keap1 protein was increased in the CD model, but the ratio of Nrf2/Keap1 was significantly lower than that in normal cells. Combined with other studies (Yao et al., 2019; Yi et al., 2020) and the characteristics of the Nrf2/Keap1 signaling pathway, it is inferred that p31-43 can cause oxidative stress in cells and activate the Nrf2/Keap1 antioxidant signaling pathway, and up-regulate the expression of Nrf2. However, p31-43 may be similar to TSN (Yao et al., 2019) in the mechanism of cell damage, both inhibiting Nrf2 activation by increasing Keap1 expression, resulting in a significantly lower Nrf2/Keap1 ratio than normal group cells, so that Nrf2 cannot effectively up-regulate the activity of antioxidant enzymes. Because Nrf2 is the upstream gene of GCLC and GCLM, the inhibition of Nrf2 expression by p31-43 will also lead to the inhibition of GCLC expression in cells, which will lead to the decrease of GSH biosynthesis rate in cells, and finally make cells unable to effectively degrade peroxidative stress, which will become more and more serious, leading to cell membrane rupture and apoptosis.

The expression of Keap1 protein with pretreatment of BSP1 was significantly decreased (Figure 8), even lower than the normal group, the expression of Nrf2 protein increased significantly and was also significantly higher than that of the normal group, leading to the significant increase of Nrf2/Keap1, and the expression of GCLC and GCLM were also significantly higher than CD model group. The expression of Keap1 protein in the BSP3 pretreatment group (Figure 8) was significantly different from that in the CD model group and was reduced to the level of the normal group, and the expression of Nrf2 was significantly up-regulated compared with the normal group, both GCLC and GCLM were significantly up-regulated. For pretreatment of BSP5 (Figure 8), compared with the CD model group, the expression of Keap1 was significantly down-regulated, while the expression of Nrf2 protein was not significantly different from that in the normal group, so the Nrf2/Keap1 value was similar to that normal group and significantly higher than CD model group. Both GCLC and GCLM were significantly up-regulated.

It was found (Yi et al., 2020) that treatment of BSPs increased the expression levels of Nrf2 mRNA and protein, inhibited the expression of Keap1, and prevented the ubiquitination and degradation of Nrf2 by Keap1. Other antioxidants can inhibit the protein expression of Keap1 by down-regulating the RNA expression of Keap1 (Lin et al., 2019), thereby releasing the inhibitory effect of Nrf2 and activating the antioxidant stress defense system. Therefore, based on the experimental results, we speculated that pretreatment of BSP1 significantly reversed the decrease in Nrf2 / Keap1 induced by p31-43, it can promote the expression of Nrf2 downstream antioxidant enzyme gene, protect the activity of the antioxidant enzyme (CAT, GR, and GST), and accelerate the efficiency of cell biosynthesis of GSH by promoting the expression of GCLC and GCLM, and relieve the oxidative stress of cells by p31-43. BSP3 has a similar effect to BSP1. pretreatment of BSP3 in the CD model also inhibits Keap1 protein expression, reduced ubiquitination of Nrf2 protein, and upregulated Nrf2 protein expression, activates downstream GCLC and GCLM genes to regulate GSH synthesis, and upregulated activity of the antioxidant enzyme (GR and GPX) to inhibit oxidative stress by p31-43. We conclude that BSP5 activates the antioxidant defense system by inhibiting or down-regulating the expression of Keap1 and thereby reducing the inhibition of Nrf2. BSP5 can promote the expression of Nrf2 in normal cells, which may be the reason why BSP5 can promote the expression of GCLC and GCLM in the CD model.

Pretreatment of BSPs can prevent oxidative damage induced by p31-43 by activating the Nrf2/keap1 signaling pathway and enhancing the activity of antioxidant enzymes in Caco-2 cells.

## 4. Conclusion

As shown in the graphical abstract (Figure 9), after the Caco-2 cells were stimulated by p31-43, they were induced to generate ROS, which led to lipid peroxidation and increased the content of MDA in cells. p31-43 also inhibits the activity of intracellular antioxidant enzymes, reduces their antioxidant capacity, and finally causes a serious imbalance of intracellular redox. At the same time, the expression of Keap1 was increased, which inhibited the expression of downstream antioxidant genes activated by Nrf2, resulting in the failure of the recovery of antioxidant enzyme activity and GSH content. And the antioxidant capacity of cells is inhibited. With the accumulation of ROS, the integrity and function of the cell membrane are damaged, and cell apoptosis or death occurs, resulting in a significant decrease in cell survival.

**Figure 9.**
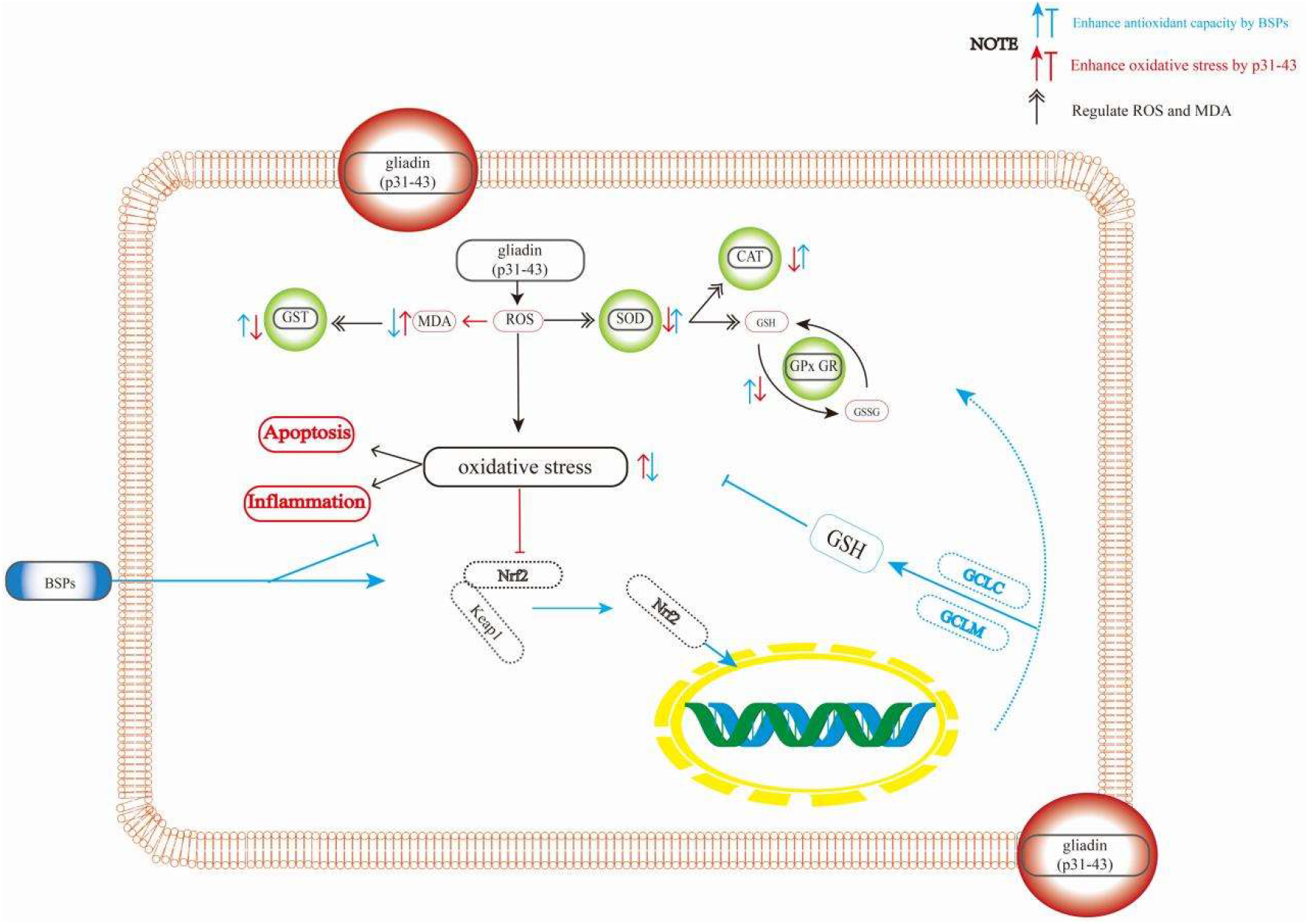
Graphical Abstract. The pattern of BSPs against oxidative damage in CD cell mode.

BSPs pretreatment can improve the antioxidant capacity of cells by increasing the activities of CAT, GR, and SOD, inhibiting the increase of Keap1 protein induced by p31-43, and activate antioxidant genes through Nrf2 protein, improving the activity of the antioxidant enzyme (CAT, GR, GPx, GST), alleviates glutathione redox cycle imbalance, promote the expression of GCLC or GCLM, improve the biosynthesis rate of GSH, thereby eliminating the content of ROS or MDA, and reduce oxidative damage.

## List of Abbreviations

CD: Celiac disease
BSPs: Black soybean peptides
h: hour
min: minute
s: second
rpm: Rounds per minute
CBS: Carbonate buffer solution
PBS: Phosphate buffer solution
DMEM: Dulbecco’s modified eagle medium
PAGE: Polyacrylamide gel electrophoresis
ROS: Reactive oxygen species
MDA: Malondialdehyde
GSH: Reduced glutathione
GSSG: Oxidized glutathione
SOD: Superoxide dismutase
CAT: Catalase
GR: Glutathione reductase
GPx: Glutathione peroxidase
GST: Glutathione s-transferase
GFD: Lifelong adherence to a gluten-free diet

## Author contributions

Conceptualization: C. C., W. N., F. W.; Data curation: F. W., C. W., X. S.; Formal analysis: C. C., E. G., W. N.; Funding Acquisition: F. W., C. W., X. S.; Investigation: F. W., C. W., N, W.; Methodology: C. C., E. G.; Project administration: F. W., C. W.; Resources: F. W., C. W.; Software: F. W., C. C., E. G.; Supervision: Q. Y., M. H., N; Validation: N. C., Y. Z.; Writing – original draf: C. C.; Writing – review & editing: F. W., N. W.

## Competing interestst

The authors declare no competing or financial interests.

